# 16S rRNA gene sequencing reveals site-specific signatures of the upper and lower airways of cystic fibrosis patients

**DOI:** 10.1101/125187

**Authors:** Sarah K. Lucas, Robert Yang, Jordan M. Dunitz, Holly C. Boyer, Ryan C. Hunter

## Abstract

**Rationale:** Chronic rhinosinusitis (CRS) is an inflammatory disorder of the sinonasal mucosa associated with microbial colonization. Metastasis of sinus microbiota into the lower airways is thought have significant implications in the development of chronic respiratory disease. However, this dynamic has not been thoroughly investigated in cystic fibrosis (CF) patients, where lower airway infections are the primary driver of patient mortality. Given the high prevalence of CRS in CF patients and the proposed infection dynamic between the upper and lower airways, a better understanding of sinus-lung continuum is warranted.

**Objective:** To compare the microbiome of matched sinus mucus and lung sputum samples from CF subjects undergoing functional endoscopic sinus surgery (FESS) for treatment of CRS.

**Methods:** Mucus was isolated from the sinuses and lungs of twelve CF patients undergoing FESS. 16S ribosomal RNA gene sequencing was then performed to compare bacterial communities of the CF lung and sinus niches. Finally, functional profiling was performed to predict bacterial metagenomes from the 16S dataset, and was used to compare pathogenic bacterial phenotypes between the upper and lower airways.

**Measurements and Main Results:** Bacterial richness was comparable between airway sites, though sinus and lung environments differed in community evenness, with the sinuses harboring a higher prevalence of dominant microorganisms. Beta diversity metrics also revealed that samples clustered more consistently by airway niche rather than by individual. Finally, predicted metagenomes showed that anaerobic metabolism was enriched in the lung environment, while genes associated with both biofilm formation and Gram identity were not variable between sites.

**Conclusions:** Sinus and lung microbiomes are distinct with respect to richness and evenness, while sinus communities have a higher incidence of a dominant taxon. Additionally, ordination analyses point to sinus and lung environments as being stronger determinants of microbial community structure than the individual patient. Finally, BugBase-predicted metagenomes revealed anaerobic phenotypes to be in higher abundance in the lung relative to the sinuses. Our findings indicate that while the paranasal sinuses and lungs may still comprise a unified airway in which lower airways are seeded by sinus microbiota, these discrete airway microenvironments harbor distinct bacterial communities.

## INTRODUCTION

Cystic fibrosis is an autosomal recessive disease caused by a genetic defect in the cystic fibrosis transmembrane regulator (CFTR) protein (1). Mutations in CFTR result in an impairment of chloride and bicarbonate transport across epithelial cell membranes, leading to impaired mucociliary clearance, a pro-inflammatory milieu, and susceptibility to bacterial infection (2). This is particularly evident in the lower airways, where chronic infections and the ensuing inflammatory response are the primary cause of CF patient mortality (3). Though *Pseudomonas aeruginosa* is the canonical pathogen of CF lung disease (4), culture-independent studies have identified remarkably complex microbiota (including fungi and viruses) that are also thought to contribute to pulmonary decline (5–9). A better understanding of respiratory microbial community dynamics and their interactions with the host will undoubtedly generate new therapeutic strategies.

In addition to the lung, CFTR defects manifest at other anatomical sites, including the biliary tree, small intestine, and the paranasal sinuses. In the latter instance, defective CFTR ion transport leads to an increase in mucus viscosity, decreased mucociliary clearance, and obstruction of the sinus ostia (10). Secondary events such as development of hypoxia and impairment of host defenses renders this niche susceptible to bacterial colonization and chronic rhinosinusitis (CRS) – a multi-symptomatic, prolonged inflammatory syndrome (11). Notably, there is a striking incidence of CRS in the CF population relative to non-CF subjects (∼16%, (12)). This is particularly evident in patients with classical CF mutations (class I-III), who, based on radiological evidence, have a CRS incidence rate of nearly 100% (13–15).

Culture-based studies have revealed that CF-associated CRS (CF-CRS) patients harbor microbiota that differ from non-CF sinus infections, including *Staphylococcus aureus* and *Haemophilus influenzae* in pediatric patients, followed by *P. aeruginosa* and other opportunistic pathogens as patients age (16). This dynamic generally follows the same temporal succession of the microbiota of the CF lung (17, 18) and several groups have demonstrated a positive correlation between the bacteriology of the upper and lower airways in CF subjects (19–21). These observations have led to suggestions that upper airway infections may spread to the lower airways (20, 21). In fact, compelling evidence has implicated the sinuses as infection foci for lung pathogens, where they first adapt to the host before descending into the lungs (22–25). Genotypic analyses of paired *P. aeruginosa* isolates between sites also suggest a direct exchange; *P. aeruginosa* cultured simultaneously from the sinuses and lungs were shown to be genetically identical in 38 of 40 subjects (95%) (26). A significant association between genotypes of isolates cultured from sinus mucus and bronchoalveolar lavage fluid has also been shown (27). These data are reinforced by studies of CF lung transplant patients, where recipient allografts were found to be re-colonized with the same *P. aeruginosa* clones as those cultured prior to transplantation (28, 29). Taken together, these observations strongly support the notion of a sinus pathogen reservoir and a single unified airway, in which the treatment of upper airway infections could have profound benefits for CF lung disease management.

While *P. aeruginosa* and S. *aureus* represent two of the most common organisms isolated from the sinuses in CF subjects (30, 31), culture-independent studies suggest that the CF sinuses are also colonized by an abundance of anaerobes (*e.g. Propionibacterium acnes*) and other non-canonical pathogens that are infrequently detected by clinical culture (30). These data are consistent with microbiome surveys of CF lung disease, which is now widely recognized to have a polymicrobial etiology (5–9). Recent sequencing studies have evaluated relationships between the microbiota of the oral, nasal and lung cavities in CF patients (32), but to our knowledge, molecular methods have not been used to assess the microbial dynamic between the CF sinuses and lungs at a community level. From a clinical perspective, these data are critical for effective therapy; at present, obtaining sinus cultures is invasive and time-consuming, making culture-guided antibiotics difficult to implement. Rather, lower airway sputum cultures are frequently used as a proxy of upper airway disease (33). Since it is generally (and perhaps incorrectly) assumed that any bacterium found in the upper airways is present in the lungs (33), antibiotics are empirically prescribed for sinus infections based on sputum culture. Patient response to this CF-CRS treatment approach is largely ineffective, and patients who fail medical management ultimately require surgical intervention (10, 34, 35). Therefore, a deeper understanding of the ecological relationships between the sinuses and lungs is needed to improve upon treatment strategies.

In this cross-sectional study, we used 16S rRNA gene sequencing to directly compare the bacterial community composition of the upper (sinus) and lower (lung) airways in a small cohort of CF patients undergoing functional endoscopic sinus surgery for the treatment of CRS. The primary objective was to compare the diversity of bacterial communities of the sinuses and lungs found by this culture independent approach. As a secondary objective, we performed predictive metagenomic profiling to assess whether differences in predicted bacterial phenotypes are reflective of the niche space in which they are found. We discuss these findings in the context of using lower airway sputum cultures to steer targeted therapies for upper airway infection.

## METHODS

### Patient cohort and specimen collection

Twelve participants with CF undergoing functional endoscopic sinus surgery (FESS) were recruited at the University of Minnesota Department of Otolaryngology, Head and Neck Surgery, and informed consent was obtained for all subjects. Prior to FESS, patients provided an expectorated lung sputum sample directly into a Sputocol sputum collection conical tube. Sinus samples were obtained from the middle meatal region under endoscopic visualization by suctioning secretions into a mucus specimen trap (Cardinal Health, Dublin, OH).

### Quantitative PCR

Bacterial burden was estimated by quantifying 16S copy number from DNA extracted from clinical specimens using qPCR. Universal 16S rRNA qPCR primers 338F and 518R were used (36, 37). QuantiTect SYBR Green (Qiagen, Valencia, CA) was used according to manufacturer’s instructions. Reactions were prepared in triplicate as described previously, with adjustments to the amplification protocol (38). Additional details can be found in the data supplement.

### DNA extraction, Library Preparation, and Sequencing

The Powersoil DNA Isolation Kit (MoBio, Carlsbad, CA) was used to extract genomic DNA from 300 μL of mucus, following the manufacturer’s protocol. Purified DNA was submitted to the UMN Genomics Center (UMGC) for 16S library preparation using a two-step PCR protocol (39). The V4 region of the 16S gene was amplified and sequenced on an Illumina MiSeq using TruSeq version 3 2x300 paired-end technology. Water and reagent control samples were also submitted for sequencing and did not pass quality control steps due to 16S rRNA gene content below detection thresholds.

### Sequence analysis

Raw 16S rRNA gene sequence data were deposited as fastq files in the NCBI Sequence Read Archive under accession number PRJNA374847. Sequence data were obtained from UMGC and analyzed using a pipeline developed by the UMN Informatics Institute in collaboration with the UMGC and the Research Informatics Solutions (RIS) group at the UMN Supercomputing Institute (40). Details are provided in the data supplement.

### Prediction of sinus and lung metagenomes based on 16S rRNA data

Metagenomes were inferred from 16S rRNA data using Phylogenetic Investigation of Communities by Reconstruction of Unobserved States (PICRUSt) (v. 1.0.0) (41). PICRUSt uses marker gene survey data to predict metagenome functional content of microorganisms through ancestral state reconstruction. We implemented PICRUSt scripts to infer metagenomes from the quality filtered OTU table. Briefly, OTUs were normalized by 16S copy number using the script normalize_copy_number.py. Normalized OTUs were used to predict KEGG orthology (KO)-based metagenomes of our samples through input into the script predict_metagenomes.py with an additional per-sample Nearest Sequenced Taxon Index (NTSI) calculation. Finally, predicted metagenomes were further categorized by KEGG pathways using the categorize_by_function.py script. Output of this script was filtered to only include those pathways that accounted for ≥1% of count data in each sample.

We then used BugBase (https://bugbase.cs.umn.edu) to summarize predicted metagenomes by bacterial phenotype. BugBase combines functionalities of PICRUSt, Integrated Microbial Genome comparative analysis system (IMG4) (42), the PATRIC bacterial bioinformatics database (43), and the KEGG database (44), to identify specific OTUs that contribute to a community-wide phenotype. The main script was run with default settings using the same filtered OTU table as used in PICRUSt.

### Statistical Analyses

All statistical analyses were carried out in GraphPad Prism 6.0 unless stated otherwise. Significance was assessed at the α = 0.05 level. A non-parametric paired t-test (Wilcoxon matched-pairs signed rank test) was used to assess significance when comparing metrics between sinus and lung samples. For alpha diversity metrics, a nonparametric t-test was used with 999 Monte Carlo permutations, implemented through the compare_alpha_diversity.py script in QIIME (45). Within-patient and between-sample type taxonomy correlations were calculated using the QIIME script compare_taxa_summaries.py using Spearman correlation with 999 permutations. Permutational analysis of variance and homogeneity of dispersion tests were carried out using the ‘adonis’, ‘betadisper’, and ‘permutest’ functions in the ‘vegan’ R package (v. 2.4.1) (46).

### Ethics Statement

Studies were approved by the Institutional Review Board at UMN (IRB no.1403M49021). Subjects provided informed written consent prior to sample collection.

## RESULTS

### Patient Cohort

The primary goal of this study was to examine variability in bacterial communities between the upper and lower airways. Twelve CF adults with CRS who were scheduled to undergo functional endoscopic sinus surgery (FESS) were recruited for the study. Prior to surgery, each patient provided an expectorated sputum sample, and sinus mucus was collected at the beginning of FESS. These samples are herein referred to as “sinus” and “lung” samples, denoting their anatomical origin. Clinical data were also collected that included CF genotype, microbiology cultures from sinus specimens, spirometry and Sino-Nasal Outcome Test (SNOT-22) scores, and prior FESS procedures (Table 1). CF genotype was available for 11 of the 12 patients, with 6 patients being homozygous for ΔF508, 3 heterozygous ΔF508, and 2 heterozygous non-ΔF508 mutations. Clinical culture data was also available for 11 of 12 sinus samples. Of these, 7 were positive for *Pseudomonas aeruginosa*, and 5 were positive for *Staphylococcus aureus*. The average SNOT-22 score across our patient cohort was 35 ± 17, consistent with previous reports of CF patients undergoing FESS (47).

**Table 1.** Summary of subject clinical data.

| Patient | Sex | Age | CFTR Genotype | FEV1% | SNOT-22 | Prior<br>FESS (#) | Polyposis | PA | SA |
| --- | --- | --- | --- | --- | --- | --- | --- | --- | --- |
| **1** | M | 25 | ΔF508/R553X | 66 | 21 |  | Yes |  |  |
| **2** | M | 31 | ΔF508/ΔF508 | 49 | 41 | Yes(2) | Yes | Yes |  |
| **3** | M | 28 | ΔF508/ΔF508 | 74 | 21 |  | Yes | Yes |  |
| **4** | M | 42 | ΔF508/ΔF508 |  |  | Yes(4) | Yes | Yes | Yes |
| **5** | M | 31 | ΔF508/ΔF508 | 95 | 20 | Yes(2) | Yes |  |  |
| **6** | M | 26 | G551D/2789+5G>A | 105 | 8 |  | Yes |  | Yes |
| **7** | F | 19 | Unknown/Unknown | 37 | 50 | Yes(4) |  |  |  |
| **8** | F | 26 | ΔF508/1717-1G>A | 98 | 59 | Yes(1) | Yes | Yes | Yes |
| **9** | F | 32 | ΔF508/ΔF508 | 31 | 34 | Yes(1) | Yes | Yes |  |
| **10** | M | 41 | 1717-1G-7A/3849+10kbc-T1 | 72 | 23 | Yes(1) | Yes |  |  |
| **11** | F | 36 | ΔF508/ΔF508 | 61 | 36 | Yes(1) |  | Yes | Yes |
| **12** | M | 31 | ΔF508/394delTT | 57 | 58 | Yes(3) | Yes | Yes | Yes |
| *Avg. +/-* |  | 30.5 +/- |  | 67.7 +/- | 35 +/- |  |  |  |  |
| *s.d.* |  | 6.5 |  | 24.3 | 17.2 |  |  |  |  |

### Bacterial load in CF sinus and lung samples

Quantitative polymerase chain reaction (qPCR) analysis of 16S ribosomal RNA (rRNA) gene copy number was performed to estimate bacterial load (Fig. 1). On average, sinus specimens contained 2.72 × 10^4^ (IQR = 2.69 × 10^3^ - 1.72 × 10^5^) 16S gene copies per 10 ng genomic DNA, while lung sputum harbored 5.33×10^5^ (IQR=6.1×10^3^ - 5.02×10^5^) per 10 ng of genomic DNA. These data suggest a modest but significant difference in 16S gene abundance between sample types (Fig. 1A, P=0.0425). Interestingly, patient age was positively associated with 16S gene abundance in sinus samples, but this relationship was not observed for the lower airways (Fig. 1B). We also assessed the relationship between bacterial load and two clinical metrics: FEV1%, the gold standard spirometry metric used to assess obstructive lung diseases (48), and the SNOT-22 survey used to assess an array of sinus disease symptoms (47). SNOT-22 scores did not significantly correlate with 16S copy number in either sinus or lung samples, nor did FEV1% (Fig. 1B). These results are consistent with previous studies where no association was found between bacterial load and lung function or quality-of-life (49). Taken together, these data suggest that the specific composition of each bacterial community rather than its overall abundance contributes to disease states in both sinus and lung niches.

**Figure 1.**
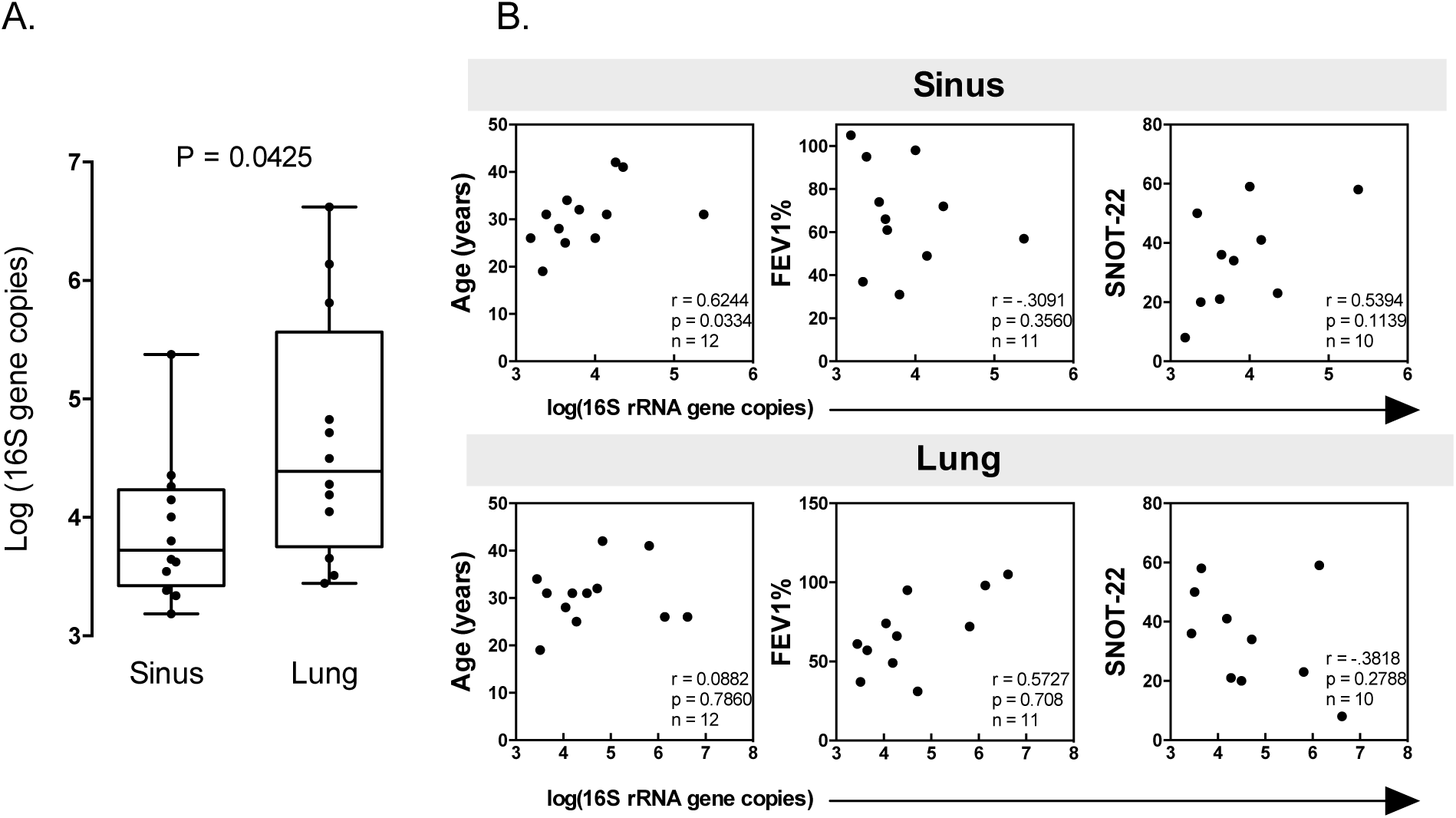
16S gene copies are greater in CF lung sputum compared to sinus samples, and correlate with patient age. Quantitative PCR was used to enumerate 16S rRNA gene copies per 10 ng of genomic DNA isolated from sinus and lung samples. **A.** Comparison of 16S rRNA gene copies in sinus and lung sample pairs by qPCR. (Wilcoxon signed-rank test *P*=0.0425). **B.** Spearman correlations with 16S rRNA gene copies and patient clinical data. In sinus samples, 16S copies are positively and significantly correlated with patient age.

### Bacterial community membership varies with respiratory tract location

Bacterial community composition of paired sinus and lung samples was profiled using 16S rRNA gene sequencing. After filtering sequences for quality and subsampling to an even depth, 51 genera were identified across all samples (Fig. 2).

**Figure 2.**
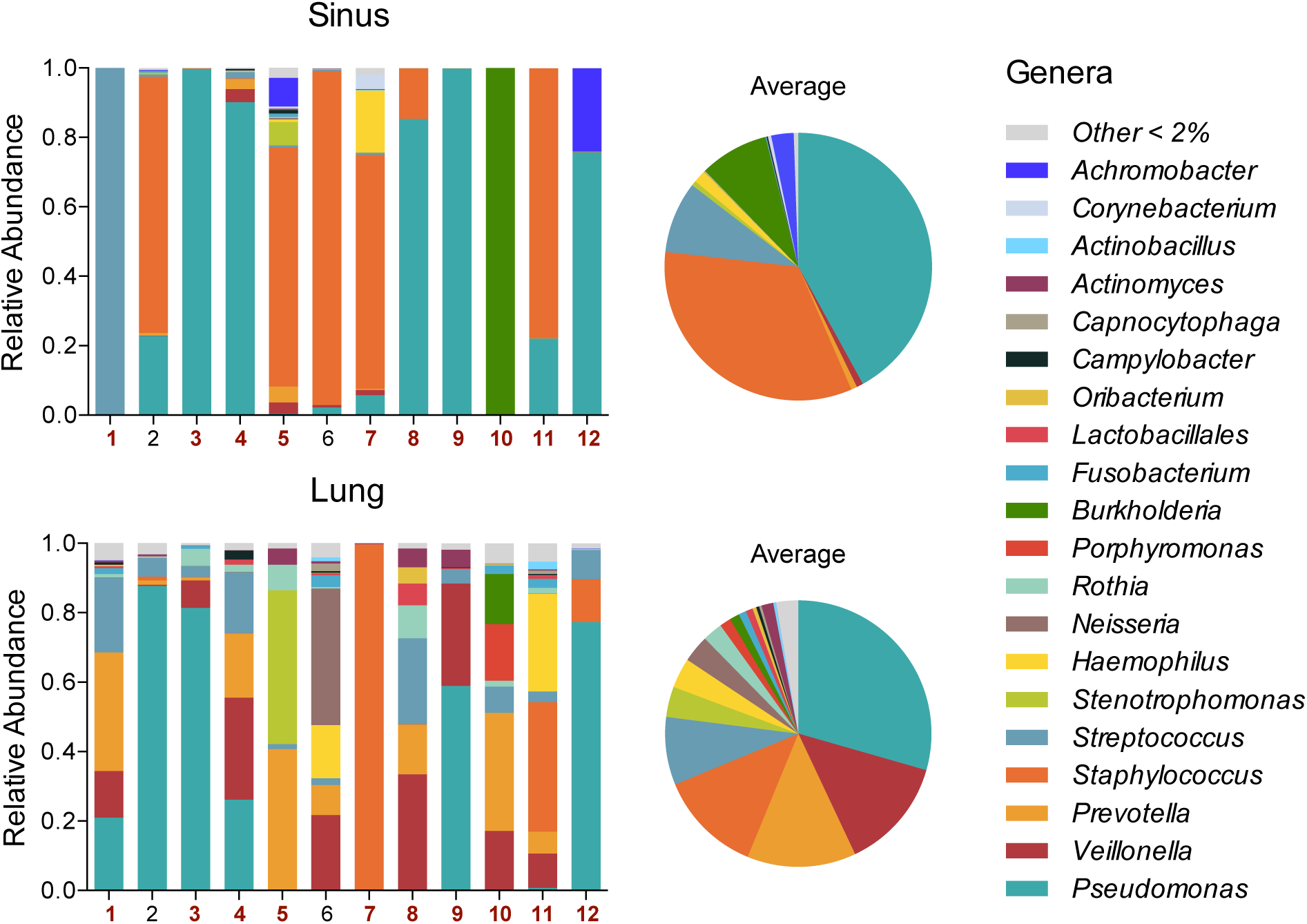
Bacterial community composition of the upper and lower airways. Stacked bar plots of relative abundances of genera in paired sinus and lung samples demonstrate dominance of *Pseudomonas, Staphylococcus*, and *Streptococcus* genera. Red patient numbers indicate samples where bacterial membership was significantly correlated within pairs. Data for these relationships is presented in Table E1. Pie charts show the average abundance of genera for each sample type across the patient cohort.

To investigate genera that accounted for the majority of sequences, we adopted the definition of a dominant genus (the most abundant genus with at least twice the abundance of the second most abundant genus) from Coburn et al. (9). A dominant genus was present in 100% of sinus samples but only 33% of lung samples. *Pseudomonas* and *Staphylococcus* were the dominant genus in five sinus samples each (42%), while *Streptococcus* and *Burkholderia* were each dominant in a single sinus sample (Fig. 2). As expected, only *Pseudomonas* and *Staphylococcus* were dominant organisms in the lower airways. The median relative abundance of the most abundant genus was 0.88 (IQR = 0.75 - 0.99) in each sinus sample, and 0.42 (IQR = 0.34 - 0.78) for lung samples. The most abundant sinus OTUs were *Pseudomonas, Staphylococcus*, and *Streptococcus*. By contrast, lung samples harbored an abundance of *Pseudomonas, Veillonella*, and *Prevotella*, consistent with previous studies (5–9). Interestingly, although many taxa were shared between sample pairs, the absence of a known CF pathogen (*e.g. Pseudomonas, Achromobacter, Staphylococcus*) in lung sputum was not predictive of its absence in the paired sinus sample, suggesting that infections at either site could be perpetuated by different pathogens within a given individual.

Based on this observation, in addition to prior studies supporting the notion of bacterial metastasis between the upper and lower airways, we were interested in the extent to which genera were shared between sites in each individual subject. Spearman correlations between genera in matched pairs revealed that within-patient similarities allowed for significant positive correlation between sites in ten of twelve sample pairs (average Spearman ρ = 0.45) (Table E1). A group-wise comparison of sinus and sputum samples showed a weaker correlation (Spearman ρ = 0.32, P = 0.001). This result highlights the potential for bacterial communities of the upper and lower airways to be similar within a given patient, yet the group correlation underlines the general dissimilarity in bacterial communities between the sinus and lung microenvironments.

### Bacterial diversity varies between the upper and lower airways

As described above, a small subset of taxa dominated most of the sequences detected in each sample. To investigate this further, two alpha diversity metrics, Observed OTUs and Shannon diversity index, were used as measures of community richness (biodiversity) and evenness (equitability) (50, 51). Observed OTUs in lung sputum were greater than in sinus samples, indicating greater taxonomic richness, though the difference was not significant (Fig. 3A). Using the Shannon diversity index, it was determined that the sinus niche was characterized by increased unevenness as compared to the sputum samples, consistent with the high prevalence of dominant genera in the sinus communities surveyed (Fig. 3B). When genera were ordered by rank, an average of 10 and 20 genera accounted for 99% of the sequences in sinus and lung samples, respectively (Fig. 3C). These data indicate that the lung environment harbors a bacterial community that is greater in both richness and evenness when compared to the sinuses in CF subjects.

**Figure 3.**
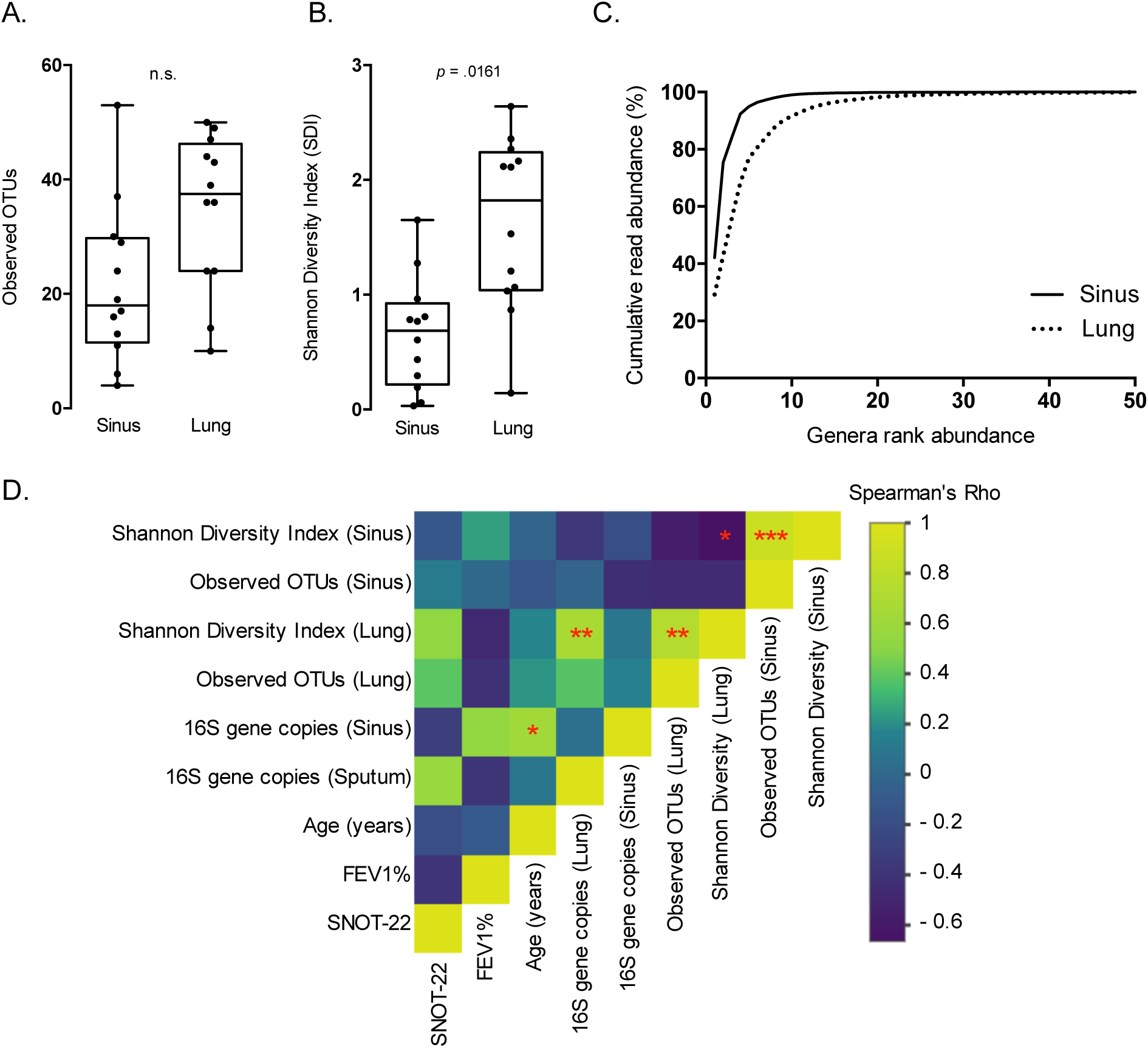
Alpha diversity of CF sinuses differs from CF lung sputum. **A.** Observed OTUs are modestly greater in lung samples compared to the sinuses. Lung samples display significantly greater evenness in OTU distribution relative to sinus samples (Wilcoxon signed rank test, *P* = 0.0161). **B.** Rank abundance curves reveal that both sinus and lung bacterial communities are dominated by a few organisms. 10 and 20 genera represent 99% of the sequences for sinus and lung samples, respectively. **C.** Spearman correlation heat map shows association of bacterial diversity (Observed OTUs, Shannon), bacterial 16S gene abundance, and clinical factors. (* P ≤ 0.05, ** P ≤ 0.01, *** P ≤ 0.001).

Spearman correlations were then calculated to assess the relationships between bacterial load, alpha diversity metrics and patient clinical data (Fig. 3D). These data revealed a significant inverse correlation between Shannon diversity in the sinus and lung (ρ=-0.664, *P* = 0.022). Because the data show a similar richness between sites, and both sinus and lung Shannon diversity revealed positive relationships with observed OTUs (sinus ρ=0.881, *P* = 3.35×10^-4^; lung, ρ=0.774, *P* = 0.007), it can be inferred that the difference in diversity is driven by evenness in these niches. Lung Shannon diversity is also positively correlated with lung 16S rRNA gene copy number (ρ=0.678, *P* =0.019). These data also reiterate the positive correlation between sinus 16S rRNA gene copy number and patient age (ρ=0.624, *P* = 0.033) as shown in Fig. 1B. Altogether, these data demonstrate that bacterial diversity differs between the sinus and lung niches in CF patients. A decrease in diversity at either site is associated with the decrease in even distribution of bacterial taxa and is associated with patient age.

### Ordination reveals respiratory tract location as a strong descriptor of phylogenetic variance

To explore multivariate relationships between samples, ordination was used to visualize the beta diversity of airway microbiota. Of interest was whether bacterial communities would cluster more closely by (a) patient or (b) sampling site. Previous studies characterizing the microbiomes of the upper and lower respiratory tract (albeit not in matched pairs) show many shared taxa between sites, but also inter-individual variation (32, 52). In addition, evidence suggests that bacterial metastasis between the upper and lower airways in CF subjects is commonplace (23, 28, 29). Based on these previous studies and our alpha diversity analyses (Fig. 2), we hypothesized that sample pairs would cluster more closely by patient, and vary with sample type (between patients).

To address this hypothesis, we compared samples utilizing the weighted Unifrac distance, an abundance-sensitive, phylogenetically relevant beta diversity metric (53). This metric was calculated to determine phylogenetic pairwise distances between each sample, then plotted using principal coordinates analysis (PCoA). Contrary to our hypothesis, variation within patient sinus and lung pairs was such that they did not cluster together nearly as strongly as they did by sample type (Fig. 4A). Permutational analysis of variance yielded a strong association with samples clustering by sample type, rejecting the null hypothesis the groupings had the same centroid (*P*=0.019). A homogeneity of dispersion test lent further confidence to this conclusion, as we could not reject the null hypothesis that the two groups of samples (sinus and lung) had the same dispersion (*P*=0.107). Lung samples demonstrated considerable phylogenetic variation when compared to sinus samples (Fig. 4A). When relative abundances from our dominant taxa (*Pseudomonas* and *Staphylococcus*) were overlaid, a strong association was revealed with much of the spatial orientation of samples in the PCoA (Fig. 4B). This analysis demonstrates that other factors, most notably the sampling site and dominant genera, contribute to the overall community structure. These analyses also suggest that the upper and lower airways select for unique bacterial community structures, despite sharing individual genera.

**Figure 4.**
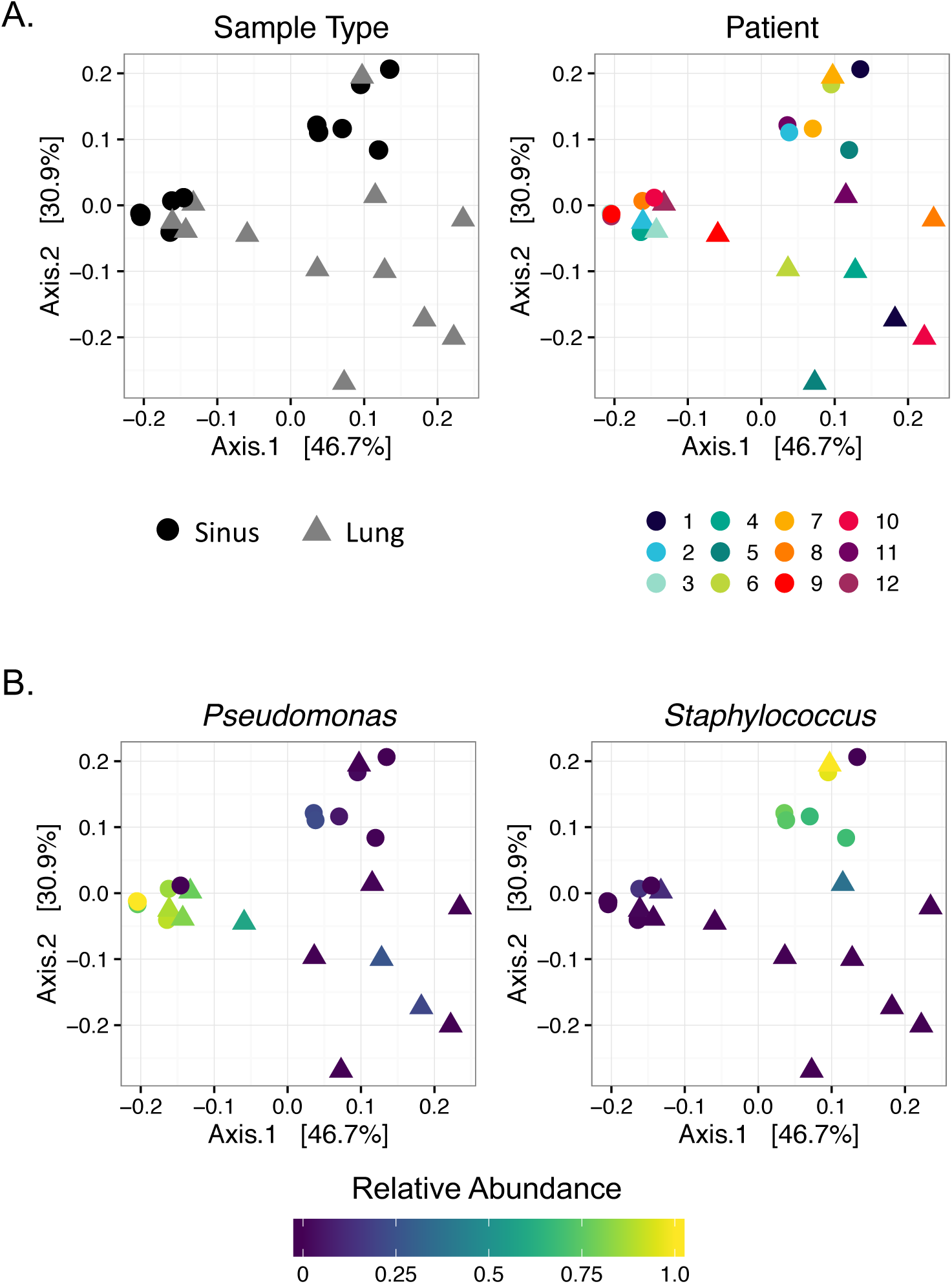
Ordination of weighted Unifrac distances shows clustering by sample type and dominant organism. **A**. Samples show more similarity by sample type rather than sampled individual (*P*=0.019). Color and shape denote patient and sample type, respectively. Sinus and lung samples do not cluster by patient, but do show clustering by sample type. **B.** PCoA colored by relative abundance of dominant organisms (defined in text), shows sinus sample grouping is highly dependent on relative abundance of *Pseudomonas* and *Staphylococcus*.

### Predicted metagenomes show conservation of most phenotypes between respiratory tract location

To gain insight into the functional capacity of airway bacterial communities, an open-source bioinformatics tool, Phylogenetic Investigation of Communities by Reconstruction of Unobserved States (PICRUSt) (41), was implemented to infer the metagenomic content of sinus and lung microbiota based on OTUs identified through 16S rRNA sequence analysis. Sequences derived from all 24 samples had a low Nearest Sequenced Taxon Index (NSTI) average of 0.017, indicating a high relatedness between bacteria found in the samples to sequenced genomes, and suggesting a high prediction accuracy for the overall dataset. Twenty unique KEGG pathways were represented in the inferred metagenomes, and showed striking similarity between sinus and lung samples (Fig. E1). This suggests that the functional capacity of the airway bacterial communities are relatively similar, despite the taxonomic diversity between sample types (Fig. 4).

To further summarize the PICRUSt output, we utilized BugBase, a bioinformatics tool that infers community-wide phenotypes from PICRUSt-predicted metagenomes, and calculates phenotypic differences between sample groups. BugBase identified that gene functions associated with an anaerobic phenotype were enriched in lung relative to sinus samples (*P* = 0.01), and could be attributed to three genera: *Veillonella, Prevotella*, and *Porphyromonas* (Fig. 5A). Gram-negative cell wall structure and the ability to form biofilms are two bacterial phenotypes often associated with pathogenicity in the human respiratory system (54). BugBase analysis shows that these bacterial phenotypes do not differ significantly between sinus and lung samples. Gram-negative bacteria contributing to this phenotype were more varied in lung samples and included *Veillonella, Prevotella, Neisseria, Stenotrophomonas* species, supporting the increased richness observed in these samples. Biofilm-forming bacteria were observed in both sinus and lung predicted metagenomes. This phenotype was influenced by the presence of *Pseudomonas* in both airway sites, but *Burkholderia* and *Achromobacter* both differentiated sinus from lung samples, while *Neisseria* and *Stenotrophomonas* were more highly represented for this phenotype in lung samples. Altogether, these data demonstrate that when bacterial phenotypes are predicted from 16S rRNA sequence data, results point towards the lung environment being a more anaerobic niche, but that other phenotypes classically linked to bacterial pathogenicity, such as biofilm formation, are similar between the two sites.

**Figure 5.**
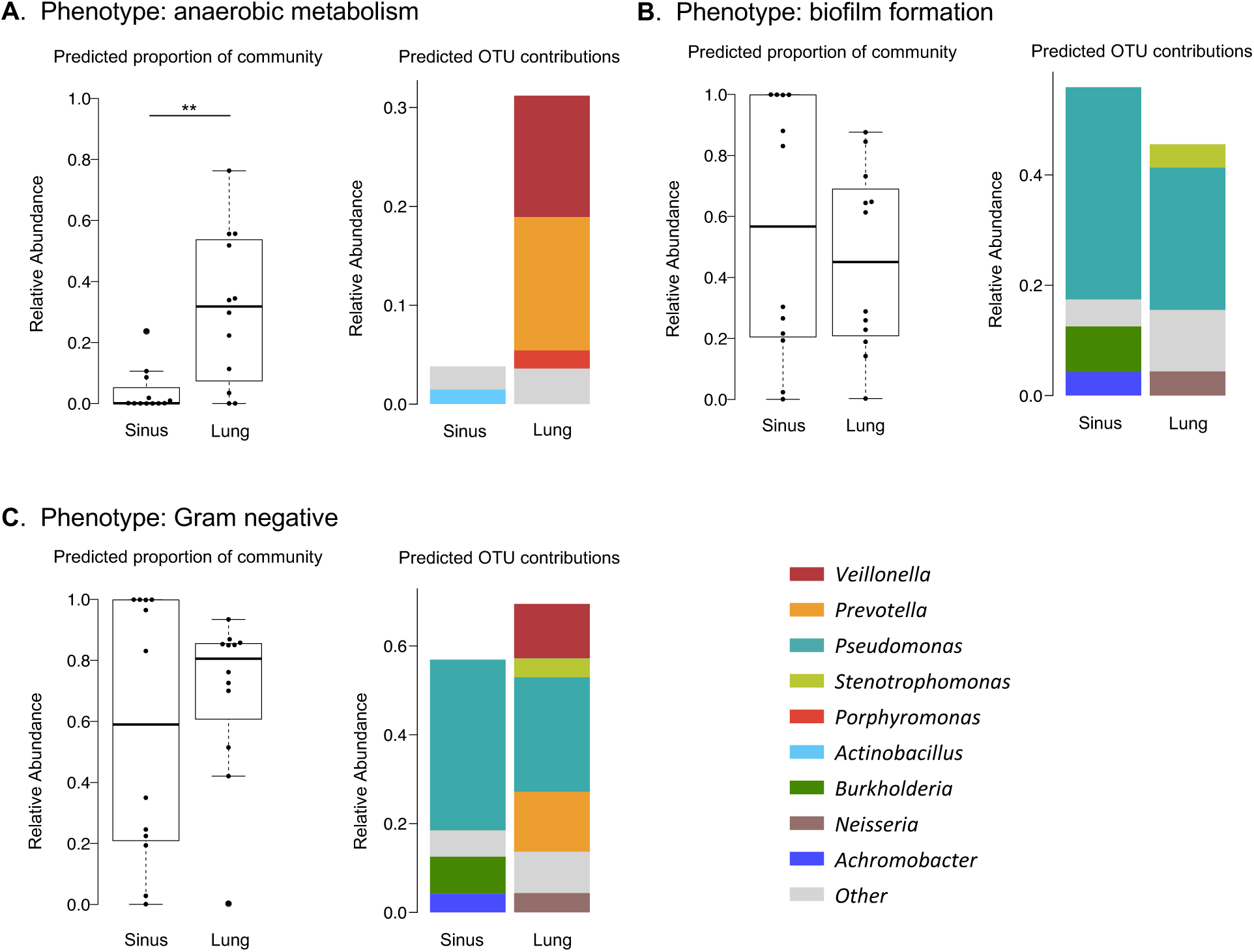
BugBase analysis of PICRUSt predicted metagenomes. **A**. Anaerobic metabolism phenotype is significantly enriched in lung samples (Wilcoxon signed-rank test, *P*=0.01). **B.** Biofilm formation does not differ significantly with sample type (Wilcoxon signed-rank test, *P*=0.57). **C.** Gram-negative phenotype is driven by presence of *Veillonella Prevotella and Pseudomonas* in lung samples (Wilcoxon signed-rank test, P=0.47). (* P ≤ 0.05, ** P ≤ 0.01, *** P ≤ 0.001

## DISCUSSION

It is poorly understood how bacterial communities of the upper airway contribute to the etiology of CF-associated CRS. It is also not known how the upper airways contribute to lower airway infections that represent the primary cause of CF patient morbidity. Numerous studies have proposed that the paranasal sinuses may be a reservoir for bacteria with pathogenic potential to adapt to the respiratory system, ultimately contributing to the worsening condition of the CF patient (23, 24, 28). It was therefore of interest to take an ecological approach towards exploring bacterial diversity in both the upper and lower airways within CF subjects with CRS, to better understand the relationship of microbiota throughout the interconnected airways.

The CF-CRS patient cohort showed striking inter-individual variability in bacterial community membership, consistent with prior microbiome studies surveying microbiota in the upper and lower airways of individuals with CF (32). Despite the considerable variability between patients, we found that while lung samples harbored an increased diversity in terms of evenness, FESS-derived sinus samples were less diverse due to the dominance of either *Staphylococcus* or *Pseudomonas*. Interestingly, this contrasts with a previous study of paired lung, nasal and throat swab samples in pediatric CF patients, which reported the inverse relationship between upper and lower airway bacterial communities (32). Given the shared taxa observed between sampling sites in our patient cohort, the data does not rule out the sinuses as a source of this colonization. Analysis of bacterial diversity in this study continually pointed to the presence of dominant and highly abundant taxa as being important in delineating these two respiratory environments. We observed that 100% of sinus communities profiled were characterized by a single dominant genus, most commonly *Pseudomonas* or *Staphylococcus. Pseudomonas* in particular is a genus associated with declining lung function in CF patients (9). It was intriguing that several sinus samples had dominant genera that are considered canonical CF pathogens, yet these bacteria went undetected in the paired lung samples. This is an important finding in the context of treatment of upper and lower airway infections, as these data indicate that respiratory cultures from the two sites are not necessarily interchangeable for the determination of antibiotic therapy.

Often in microbiome surveys, individual community members may differ dramatically from sample to sample, but functional capabilities remain conserved (55). The present study highlights this relationship in the airways. Contrasting the differences in bacterial diversity between the sinus and lung niches, we found that the predicted functional capacity of the bacterial communities in both niches was similar. Yet, interestingly, a small number of BugBase-inferred phenotypes differed significantly between sample groups. In the context of the bacterial contribution to CF disease, it is plausible that there are many similarities in the microenvironments of the upper and lower respiratory system that may contribute, or even result from these conserved microbial functions. These data support the hypothesis that the sinuses can harbor bacterial genera, such as *Pseudomonas*, that are well adapted to the respiratory environment and contribute to lower lung morbidity.

Altogether, data presented here demonstrate that despite the unified nature of the airways, bacterial communities of the sinuses and lungs are distinct. Whereas CF sinuses are typically dominated by a single organism, the lower airways exhibit greater diversity, marked by the presence of an anaerobic bacterial phenotype. Despite differences in diversity, bacterial populations share many predicted functional capabilities. Shared taxa between the sample pairs reflects the interconnectedness of the airway, though differences suggest CF sinus and lung microenvironments may play a crucial role in dictating the prevalence and abundance of canonical CF pathogens in the lower airways. Data presented here can be translated to the clinic by informing caregivers to utilize respiratory cultures originating from samples taken from the location of infection. Secondly, we advocate for the usefulness of 16S rRNA gene sequences in the clinical setting to supplement clinical culture data. In the future, it will be imperative to conduct longitudinal experiments surveying paired upper and lower respiratory tract samples in CF patients to explore temporal shifts in bacterial diversity as airway infections evolve.

## ACKNOWLEDGEMENTS

We acknowledge the Minnesota Supercomputing Institute (MSI), Daryl Gohl and John Garbe at the UMN Genomics Center for sequencing assistance. We thank Ali Stockness and Rebecca Dove of the Department of Otolaryngology, and the members of BioNet at the University of Minnesota for facilitating sample collection. Our thanks go out to Tonya Ward, Dan Knights, and the Knights Lab at UMN for their assistance with the BugBase analysis.

